# Chemotactic responses of *Trypanosoma brucei* procyclic forms to proline and other metabolites

**DOI:** 10.64898/2026.07.02.736028

**Authors:** Sebastian Knüsel, Mattias Benninger, David M. Versluis, Robert Insall, Calvin Tiengwe, Isabel Roditi

**Author notes:** Equal contributions (Mattias Benninger, David M. Versluis).

## Abstract

Many protozoan parasites have complex life cycles entailing migration through different organs in their hosts, but the cues guiding them remain poorly understood. Using a semi-solid plate motility assay, we show that early procyclic forms of *Trypanosoma brucei*, the first stage to develop in the tsetse fly midgut, perceive several metabolites - including glucose, glycerol and proline - as chemoattractants, while the glycolytic end-product succinate acts as a repellent. During adaptation in the fly, *T. brucei* switches from glucose/glycerol to proline as its primary energy source. We show that the parasite’s chemotactic response towards proline requires adenylate cyclase ACP5 and the cyclic AMP response protein CARP3, two components of signalling pathway involved in pH sensing. These results further support a role for *T. brucei*’s expanded repertoire of receptor adenylate cyclases as environmental sensors that guide navigation through the host.

**Highlights:**

- The plate assay reported here can be used to study how trypanosomes sense external signals and migrate in response to them.
- Early procyclic forms of *T. brucei* respond to a variety of metabolites and exhibit different patterns of collective migration on plates, depending on the stimulus.
- The response to proline operates through cAMP signalling and requires ACP5 and CARP3, two components also required for pH sensing.

---

Chemotactic responses in nature are widespread and varied. These range from individual bacteria seeking nutrients to the collective migration of neural crest cells during mammalian embryogenesis, neutrophils homing in on pathogens or metastatic cancer cells migrating between organs. In some cases, cells are reacting purely to external gradients, but in others they create or influence the gradients that guide them in the right direction, a phenomenon known as self-steering (1,2). While these different systems have been studied in depth, there is a knowledge vacuum when it comes to parasitic protozoa. Parasites with complex life cycles often alternate cycles of proliferation and differentiation, migrating between different host organs/tissue types in the process. Surprisingly, it is rarely asked how the parasites orient themselves within their hosts and what enables them to home into specific organs.

*Trypanosoma brucei* ssp. are two-host parasites causing human sleeping sickness and the cattle disease Nagana. Their definitive host is the tsetse fly, while mammals are intermediate hosts. Trypanosomes lack G-protein coupled receptors and heterotrimeric G proteins, which act as sensors and signal transducers in yeast and multicellular eukaryotes.

Instead, they encode an unusually large number of receptor adenylate cyclases which have been proposed to exercise a similar function (3). ACP1 - ACP6 are six adenylate cyclases that were characterised in procyclic (insect midgut) forms (4,5). ACP2 is distributed along the length of the flagellum, while the other five ACPs are concentrated at the flagellar tip (4–6).

We previously demonstrated that the early procyclic form, the first life-cycle stage to develop in the tsetse fly, can both sense and create pH gradients when cultured on agarose plates containing glucose (6). The cells initially form a colony as they multiply at the inoculation site. Glucose metabolism results in a local pH gradient in which the more acidic centre of the colony acts as a repellent, and the more basic environment outside the colony acts as an attractant. This bimodal form of pH sensing, which was initially termed social motility (SoMo) (7) leads to the formation of discrete projections containing thousands of cells. Early procyclic forms also respond to exogenous acid or alkali spotted outside the colony, with the projections orienting away from the acidic region and towards the alkaline region on the plate (6). In contrast to the early procyclic form, the late procyclic form, the next life-cycle stage in the fly, is not repelled by acid, but is still attracted to alkali (6). The two forms can be distinguished by the expression of the surface marker GPEET procyclin in early procyclic forms and its absence in late procyclic forms (8).

The formation of projections and pH sensing are dependent on cyclic AMP signalling (4,6,9). Mutants lacking phosphodiesterase B1 (PDEB1) or cAMP response protein 3 (CARP3), or a single knockout of ACP5 do not form projections on plates in the presence of glucose (6,10). As shown previously, this is not due to impaired motility or growth (6,10). PDEB1 and CARP3 null mutants also show defects in migration through tsetse tissues (6,10,11). The single knockout of ACP5 was not tested *in vivo*. In contrast to ACP5, which is required for the formation of projections during SoMo, ACP1 and ACP6 seem to exert an inhibitory effect.

Depletion of ACP1 or ACP6 by RNAi (4) or deletion of one copy of ACP1 or both copies of ACP6 (6) lead to the phenomenon of hyper-SoMo, in which the colonies produce greater numbers of protrusions and do so more rapidly than wild-type cells.

N-acetylglucosamine binds to trypanosome hexose transporters, competing with glucose uptake (12). Early procyclic forms cultured on plates lacking glucose (SDM80 with 10% foetal bovine serum, supplemented with 6 mM N-acetyl glucosamine to block uptake of serum glucose), do not spontaneously form projections for the first 5 days as they do not acidify their environment (6). This allowed us to test a variety of compounds for chemotactic effects on *T. b. brucei* 427 Bern. Cultures of this stock are stably >95% GPEET-positive in the presence or absence of glucose (6) meaning that the majority are early procyclic forms. Figure 1 shows the response to different metabolites. For this assay, 2 x 10^5^ cells were inoculated at the centre of the plate and incubated for 5 days; this was followed by spotting test compounds at a fixed distance on either side of the colony and incubating the plates for a further 3 days. The growth of control colonies for eight days, without spotting a test compound at day 5 (Figure 1, upper panel, right and lower panel, second from right), is reminiscent of classic Adler rings created by bacteria, as nutrients are consumed at the boundary and the wave of cells at the front moves towards higher concentrations. Malate, pyruvate, proline, mannose, glucose, glycerol and fumarate acted as chemoattractants. The response to malate or pyruvate overlays the Adler ring with a typical chemotactic response. The migration towards proline is also chemotactic, as well as appearing to stimulate cell growth closer to the source. The projections moving towards proline do not splay apart. The other attractants produce a response in which cells advance in discrete, finger-like projections that are strikingly reminiscent of reaction-diffusion responses, in which chemotaxing cells release a repellent and the final pattern reflects the interaction between the two. The migration towards mannose and glucose is likely to be due to self-generated pH gradients resulting from glycolysis, in the same way that this occurs on plates containing glucose throughout. In this case, however, high local concentrations of glucose or mannose, diffusing outwards from where they were spotted, would out-compete N-acetylglucosamine and be taken up by the cells. Acidification would then cause repulsion between projections, and result in the fingering pattern. Glycerol bypasses the hexose transporters and feeds into glycolysis, and so will cause strong pH gradients, which may explain why it leads to even more distinct protrusions than glucose and mannose. We do not know if a repellent is produced in the presence of fumarate, but the pattern of chemotaxis implies it. Succinate, which is one of the excreted end-products of glucose metabolism by procyclic forms, was the only compound that acted as a repellent. This was not a pH effect, as the solution was adjusted to ∼pH7.4. The other excreted metabolite, acetate, had no effect at this pH.

**Figure 1.**
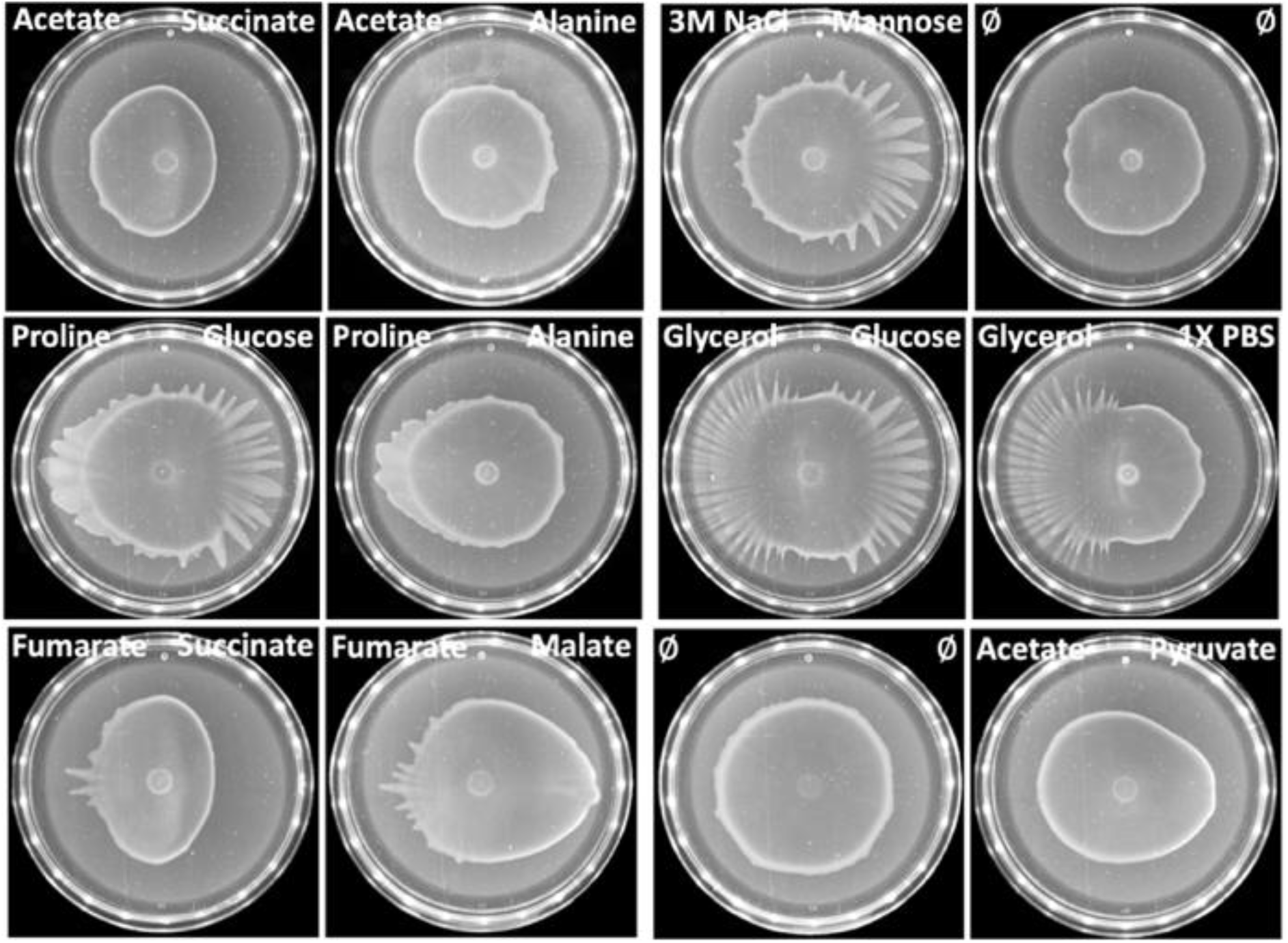
Early procyclic forms respond to different compounds. *T. b. brucei* 427 Bern was adapted to glucose-free medium by passage for at least 7 days in SDM80 supplemented with 10% FBS and 6 mM N-acetylglucosamine. The cells used in this figure were cultured in this medium for 30-37 days. Cells were pelleted and resuspended at a density of 4 x 10^7^ ml^-1^. Five µl (2 x10^5^ cells) were inoculated on the surface of plates containing the same medium and supplements (6). After 5 days incubation, 10µl of a 1M solution of a test compound (adjusted to pH∼7.4, except for pyruvate (pH∼7.0) and 3M NaCl (pH∼5.5)) was spotted on the left or the right side, 13 mm away from the edge of the colony. The plates were photographed 66-70 hours later. Two plates (upper panel, right and lower panel, second from right) are controls where nothing was spotted. 3M NaCl was tested as a control for 1M solutions of sodium salts of trivalent anions (not included in this study).

## PBS

phosphate-buffered saline; ϕ: no addition.

## Methods

Cell lines were adapted from SDM79 (19) to SDM80 medium, which is a derivative lacking glucose. Addition of 5 mM glucose to SDM80 gives it the same composition as SDM79.

Home-made SDM80 was used for Figure 1; SDM80 was sourced from Life Technologies Ltd (Paisley, U.K.) for the experiments shown in Figure 2. Plates were produced as described in detail by Imhof et al. and Shaw et al. (6,20). Following inoculation, plates were incubated in a single layer at 27°C, 2.5% CO _2_ (cells facing upwards) for the times indicated. All cell lines used here have been described previously (6). The CARP3 addback expresses a haemagglutinin-tagged version of the protein. The ectopic copy of the gene is integrated into a procyclin locus and expressed from a procyclin promoter.

**Figure 2.**
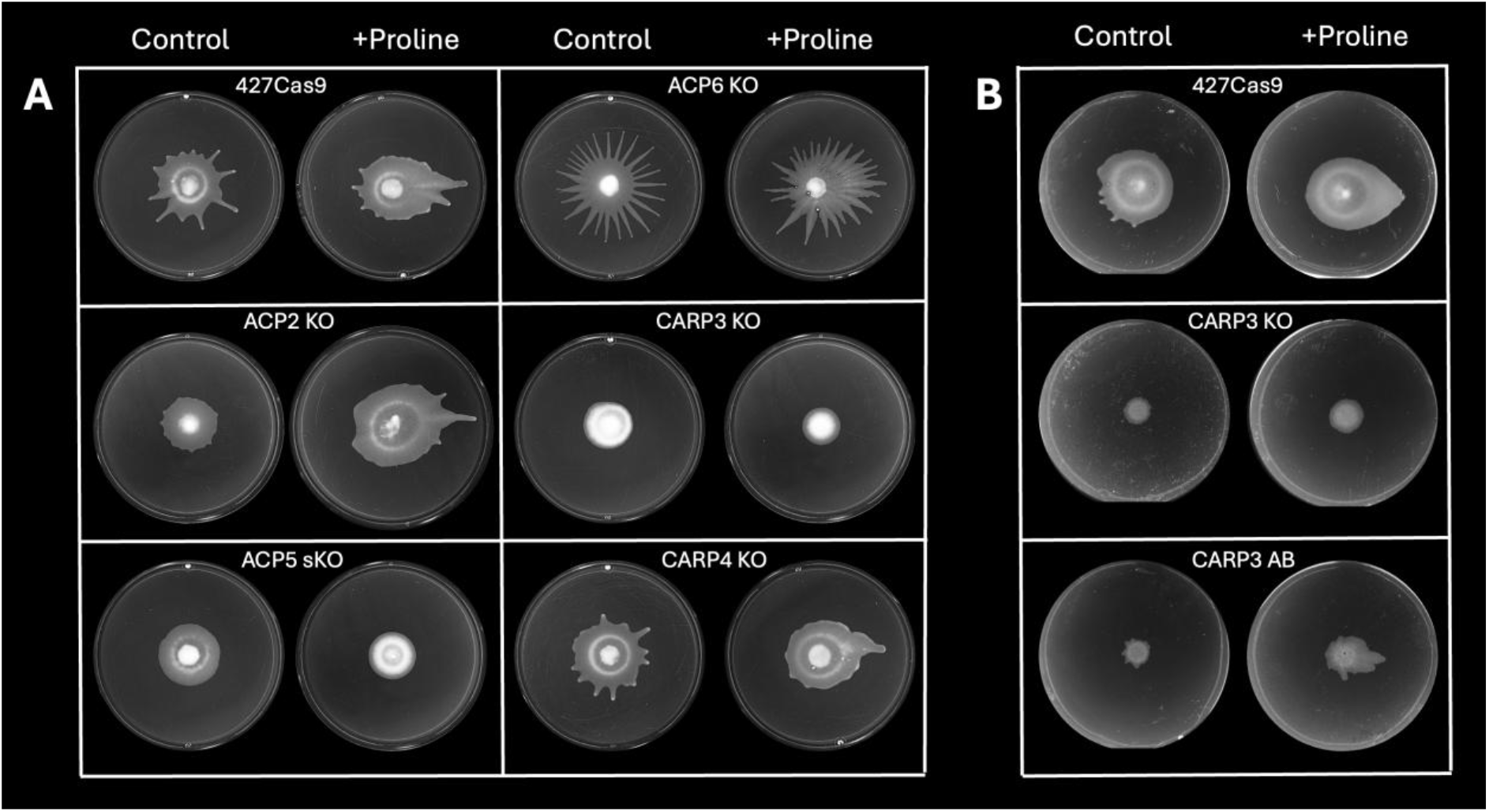
Cyclic AMP signalling is required for the response to proline. **A:** The parental line 427Cas9, ACP5 single knockout (sKO) and double knockouts (KO) of ACP2, ACP6, CARP3 and CARP4 were tested for their response to proline. Cells grown in SDM80 supplemented with 5 mM glucose and 10% FBS were pelleted and most of the supernatant removed. Cells were resuspended in the residual medium at a density of 4 x 10^7^ ml^-1^. Five microlitres (2 x10^5^ cells) were inoculated onto the surface of plates containing SDM80, 10% FBS and 6 mM N-acetylglucosamine. Controls: plates were incubated for 8 days without spotting any compound; + Proline: plates were incubated for 5 days, after which 10µl of a 1M solution of proline (pH∼7.4) was spotted on the right side, 13 mm away from the edge of the colony. Plates were photographed 72 hours later. **B:** The parental line 427Cas9, CARP3 knockout (CARP3 KO) and the CARP 3 addback (CARP3 AB) were adapted to growth in SDM80 supplemented with 10% FBS and 6 mM N-acetylglucosamine for at least 7 days before inoculation onto plates containing the same medium, as described for Figure 1. Plates were treated as described for Figure 2A and photographed 70-72 hours after application of proline.

When trypanosomes are ingested by a tsetse fly, they transition from using glucose and/or glycerol as energy sources to a reliance on proline uptake and catabolism (13–15). To test whether cAMP signalling was involved in the perception of proline, we tested a derivative of *T. b. brucei* 427 Bern that stably expressed Cas9 and several knockouts in this background (6). For this preliminary screen, cells were cultured in SDM80 + 6 mM glucose (including 1 mM glucose contributed by the foetal bovine serum), then inoculated directly on plates containing SDM80 + 6 mM N-acetylglucosamine (Figure 2A). These cultures had an intermediate SoMo phenotype akin to cultures containing 1 mM glucose (6), forming short projections over a period of 8 days, presumably because of the residual glucose transferred with the cells. In addition to the 427Cas9 parental line, the ACP2, ACP6 and CARP4 double knockouts all exhibited collective migration towards proline, indicating that these genes were not essential for the response (Figure 2A). By contrast, the ACP5 single knockout and the CARP3 double knockout did not migrate towards proline. In this respect, the responses to acid sensing and proline overlap. All cultures were still >97 % GPEET-positive, which excludes the possibility that the CARP3 knockout and the ACP5 single knockout had become non-responsive through differentiation to late procyclic forms. Finally, we tested the responses of the 427Cas9 parental line, the CARP3 knockout and CARP3 AB, a haemagglutinin-tagged addback derived from the knockout, to proline. For these assays, the cells were first adapted to SDM80 + 6 mM N-acetyl glucosamine in liquid culture for at least 7 days before plating (Figure 2B). CARP3 AB has previously been shown to restore the response to exogenous acid (the knockout was still able to sense exogenous alkali, suggesting that this part of the pH response can bypass CARP3) (6). CARP3 AB also responded to proline, with collective migration towards it (Figure 2B). Although the phenotype was not rescued completely, this is not unusual in trypanosomes. If an addback is expressed ectopically, it will not be expressed at the same level and will not be subject to the same controls as the endogenous gene.

In conclusion, early procyclic forms of *T. brucei* are responsive to a variety of extracellular signals and exhibit different patterns of collective migration on plates, depending on the stimulus. The chemotactic response to proline operates through cAMP signalling and requires ACP5 and CARP3, the same two components required for pH taxis. Since CARP3 associates with a variety of adenylate cyclases at different stages of the trypanosome life cycle (6,11), it may operate as a node that translates sensing of different extracellular stimuli into motion along gradients. The assay we have established can readily be implemented to study how trypanosomes sense and migrate in response to other signals. The mutants tested here also give a first indication of the sensing mechanism. The ability of adenylate cyclases to sense diverse external stimuli is likely to be combinatorial, since trypanosome adenylate cyclases are activated by dimerisation (16). Although it was not possible to generate a ACP3 knockout to test its effect on pH sensing (6), we showed previously that ACP5 coprecipitates with ACP3 and vice versa, suggesting that they operate as a heterodimer (6). Since *T. brucei* encodes approximately 70 adenylate cyclases, many of which are differentially expressed (14), there could be several thousand permutations with different specificities and sensitivities during the course of the life cycle. In addition to metabolites, early procyclic forms can sense exosomes (17) and live bacteria (18), but it is not known how these are perceived. Just as the inputs are likely to be combinatorial, the outputs may be as well. In their natural environment of the tsetse digestive tract, trypanosomes will encounter many signals at once: some from the fly, others from the fly microbiota, and some self-generated through metabolic activity. The direction and form of collective migration are then expected to reflect the integration of overlaid attractant and repellent gradients rather than any single cue. The plate assay described here offers a route to predicting which combination of metabolites attract, repel or modulate chemotaxis, ultimately providing insights into how trypanosomes navigate through their hosts.

## Acknowledgements

We thank Dan Davis (Head of Department, Life Sciences, Imperial College London) for granting visiting scientist status to M.B. and D.M.V., enabling the establishment of SoMo plate assays and experimental work at Imperial College. We are grateful to Eva Gluenz for facilitating M.B.’s visit to Imperial College, Ruth Etzensperger for culturing and shipping mutant stocks, and Sebastian Shaw for advice.

